# Repetitive mild closed-head injury induced synapse loss and increased local BOLD-fMRI signal homogeneity

**DOI:** 10.1101/2024.05.24.595651

**Authors:** Marija Markicevic, Francesca Mandino, Takuya Toyonaga, Zhengxin Cai, Arman Fesharaki-Zadeh, Xilin Shen, Stephen M. Strittmatter, Evelyn Lake

## Abstract

Repeated mild head injuries due to sports, or domestic violence and military service are increasingly linked to debilitating symptoms in the long term. Although symptoms may take decades to manifest, potentially treatable neurobiological alterations must begin shortly after injury. Better means to diagnose and treat traumatic brain injuries, requires an improved understanding of the mechanisms underlying progression and means through which they can be measured. Here, we employ a repetitive mild closed-head injury (rmTBI) and chronic variable stress (CVS) mouse model to investigate emergent structural and functional brain abnormalities. Brain imaging is achieved with [^18^F]SynVesT-1 positron emission tomography, with the synaptic vesicle glycoprotein 2A ligand marking synapse density and BOLD (blood-oxygen-level-dependent) functional magnetic resonance imaging (fMRI). Animals were scanned six weeks after concluding rmTBI/Stress procedures. Injured mice showed widespread *decreases* in synaptic density coupled with an i*ncrease* in local BOLD-fMRI synchrony detected as regional homogeneity. Injury-affected regions with *higher* synapse density showed a *greater increase* in fMRI regional homogeneity. Taken together, these observations may reflect compensatory mechanisms following injury. Multimodal studies are needed to provide deeper insights into these observations.

## Introduction

Repetitive mild traumatic brain injury (rmTBI), sustained through sport, military service or domestic violence, is associated with a myriad of long-term symptoms including impaired cognition, problems with impulse control, aggression, dementia, and suicide (see reviews, Ramalho & Castillo et al. 2015; Ozga et al. 2018; Maresca et al. 2023; Chen et al. 2022)^1–4^. Symptoms can take a decade or more to manifest. Yet, the neuropathological processes which give rise to deleterious long-term outcomes must begin shortly after injury. This latency makes the management of patients with rmTBIs especially challenging. Research efforts are needed to uncover methods for assessing rmTBI outcomes and the efficacy of novel intervention strategies. To this end, comprehensive multimodal approaches using animal models are essential for identifying the neuropathological changes which contribute to neurodegeneration after head injury that are accessible by clinically applicable modalities.

Modeling the biophysical and mechanical forces experienced during rmTBI by humans poses challenges in animal studies. Consequently, a number of models have been developed^5–10^ which, to varying degrees, recapitulate various aspects of rmTBI pathology^5, 9^. Given the significance of pathological tau species (herein, p-tau) in some human TBIs^11^, it is imperative for animal models to explore recapitulating this phenotype. Here, we use a mouse model which combines rmTBI, using a closed-head injury (CHI), and stress using a chronic variable stress paradigm (CVS), referred to as rmTBI/Stress henceforth^12, 13^. Critically, this combination results in persistent neurocognitive and behavioral deficits, neuroinflammation, and p-tau (which we and others have shown previously, see Tang et al. 2020; Wen et al. 2019; Ojo et al. 2014; Fesharaki-Zadeh et al. 2020)^12, 14–16^. While this may not be of direct relevance to injuries sustained through sport, it is likely of high relevance to veterans, and victims of assault and domestic violence. Elevated p-tau in particular has been linked to more severe long-term outcomes^11, 17–21^. Furthermore, CHI does not involve a craniotomy, which enables the inclusion of multiple impacts, and is, overall, a better model of the injuries sustained by humans^22, 23^.

Typically, studies of rmTBI in animal models use a battery of behavioral assessments and/or immunohistological (IHC) measurements^9^ to characterize injury and/or recovery. Here, we use the same model as our group’s previous work^24^, where it was shown that the microlesion at the impact site is restricted to superficial cortical layers, with the perilesional and distal sites remaining unaltered by routine histological stains. The neuronal marker NeuN revealed limited or no decline in neuronal density relative to sham controls. In the present study, we elect to focus on *in vivo* assessments of injury as these are more readily translatable to patient populations. The use of *in vivo* imaging modalities, such as magnetic resonance imaging (MRI) and positron emission tomography (PET), in animal models is not as widespread as is behavior and IHC studies, despite their use within the patient population.

Functional MRI analyses from human rmTBI patient studies, such as measures of regional homogeneity (ReHo), at various times after injury, show markedly heterogeneous results in terms of the brain regions and circuits affected^25–29^. Similarly, heterogeneous alterations in fMRI functional connectivity (FC), have been reported^30–34^. Although significantly less studied, animal models of rmTBI, at various times after injury (either single or multiple concussions) also report diverse alterations in FC and ReHo^35–38^. Combining fMRI with another clinically accessible modality, such as PET, may provide further insights into heterogeneous fMRI changes observed across rmTBI animal models.

Combining advanced neuroimaging modalities, such as PET and fMRI, may yield biomarkers that can depict tissue status, structural and functional brain alterations, ultimately aiding in diagnosis of the complex heterogeneous disease process of TBI. Improved characterization of post-injury mechanisms stands to improve rmTBI diagnosis, classification, and prognosis. In an effort to establish increased translational relevance of preclinical studies, we employ two clinically accessible modalities - PET with the synaptic vesicle glycoprotein 2A ligand marking synapse density and fMRI - to assess abnormalities in a mouse model where rmTBI and CVS are combined. To the best of our knowledge, no studies report using the synaptic vesicle glycoprotein 2A (SV2A) marker to assess changes in synaptic density in either humans, or animal models of TBI. Moreover, there are no studies that combine this measure with fMRI.

Our results show widespread decreases in synaptic density (PET) and detect an increase in ReHo (fMRI) in brain regions with decreased synaptic density. SV2A-PET standardized uptake value ratios (SUVRs) and BOLD (blood-oxygen-level-dependent) fMRI ReHo show a positive correlation in healthy mice (across the whole brain). With injury, this relationship (SV2A-PET SUVRs vs. BOLD-fMRI ReHo) strengthens. Moreover, separate from the impact foci and in areas without any physical damage (based on structural MRI), the affected brain regions in injured mice with higher synaptic density show a greater proportional increase in ReHo than do regions with lower synapse density. Functional connectivity changes between regions with decreased synapse density are also reported. Taken together, these observations may reflect partial spontaneous neural circuit compensation for injuries sustained. Further multimodal studies are needed to provide clarification and deeper insight into these observations.

### Methodology

Our study exclusively examined male mice. Thus, due to the known differences between males and females in TBI (see review, Biegon 2021)^39^, it is unknown whether the findings presented here are relevant for female mice. All experiments and procedures are conducted following the Institutional animal care and use committee (IACUC) approved protocols at Yale University. Wildtype male C57BL/6J mice (N=20) from Jackson Labs (stock number: 000664) at 16 weeks of age are used. All mice are kept in standard housing under a 12h light/dark cycle with food and water provided *ad libitum* throughout the experiment (with the exception of overnight food deprivation which was one of the possible stress exposures included in our CVS paradigm, see below).

### Repetitive mild closed-head injury combined with chronic variable stress

We combine rmTBI, using a CHI, and CVS (stress), referred to as rmTBI/Stress hereafter^12^. CVS is induced as described previously (N=10)^12^. Briefly, CVS consists of 2-weeks of daily exposure to 3/5 aversive stimuli: (1) 3- minute cold water swim (16-18°C), (2) overnight food deprivation with access to water *ad libitum*, (3) 3-hours in a cage with 300ml of water added (4) 3-hour exposure to a cage tilted at a 45° angle, and (5) 15-minutes immobilization in a flat-bottomed restraint chamber (Braintree Scientific Instruments). Aversive stimuli are randomized in each session for each mouse to simulate the unpredictable nature of psychological trauma, and limit habituation. Sham-stress mice are transferred to a single occupancy cage for the duration of the stressor. Daily rmTBI occurs within 1-3 hours of stress exposure.

Prior to each injury, anesthesia was induced with 3.5% isoflurane (in O_2_, 1.0 L/min) and maintained at 3% until immediately after impact when anesthesia is discontinued (< 10 minutes). The head is shaved, the animal placed in a stereotaxic frame (**Figure 1**), and rmTBI induced using a 5mm diameter tip operated by an electromagnetic impactor (Leica Microsystems, Buffalo Grove, IL). The impactor tip is placed 5mm lateral of the sagittal line, 5mm caudal of the eye, and at 20° from vertical (**Figure 1**). The impact is at a velocity of 5m/sec, depth 1mm and dwell time 100ms. Impacts alternate between the right and left hemisphere on consecutive days to minimize the probability of hemorrhage.

**Figure 1:**
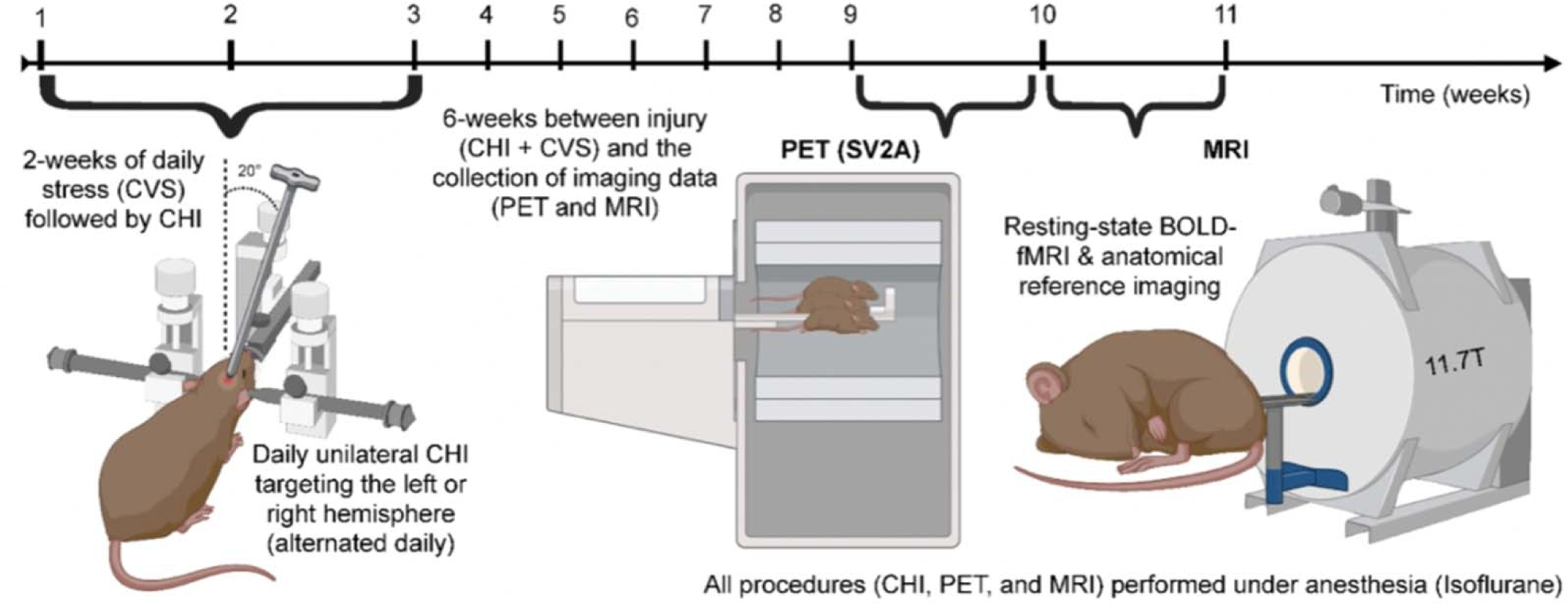
Timeline. Mice (N=10) are subjected to daily random stressors for 2 weeks (chronic variable stress, CVS), followed immediately by a mild brain injury (repeated daily) over a two-week period. Six-weeks after concluding rmTBI/Stress procedures, PET data and MRI data are collected (one-week apart). For all methodological details (e.g., sequences used, anesthesia, etc.), see **Methods**.

### AZD0530 treatment

A subset of 5 injured mice receive AZD0530 treatment (saracatinib), while the rest are given the vehicle. Amongst tau-interacting proteins is the neuronally-enriched cytoplasmic tyrosine kinase, Fyn, a member of the Src family. In preclinical models of Amyloid-ß-triggered deficits, Fyn inhibition with the orally available kinase inhibitor AZD0530 (saracatinib) reduces synaptic loss, memory deficits and p-tau ^40^. Mice began treatment (or vehicle) 24-hours following the last day of rmTBI/Stress^12^. Treatment doses of 5mg/kg/day are given in two equal aliquots by oral gavage until the study endpoint. The vehicle is 0.5% weight/volume (wt/vol) hydroxypropyl-methylcellulose per 0.1% wt/vol polysorbate 80. Each dose is 250µl. Treatment effects are not detectable (possibly due to a subtherapeutic low dosage). All 10 injured mice (regardless of treatment-status) are grouped together in our analyses (but figures include shaded dots to indicate treated vs. vehicle). Independent two-tailed t-tests are used to compare the synaptic density changes between treated and untreated animals for a number of different brain regions (**Figure S2**).

### Control group

N=3 mice undergo sham-rmTBI and sham-stress. Sham-stress mice are transferred to a single occupancy cage for the duration of the stressor. Sham-CHI mice are anesthetized and have their heads shaved but do not undergo impaction. To address this small sample size, N=7 mice are added which underwent identical imaging procedures but did not undergo sham-CHI or sham-CVS. To assess the similarity of sham and naïve mouse datasets, we compare whole-brain FC between each mouse and the group average using Spearman’s rho (**Figure S1**). No notable deviations are found. Thus, all 10 uninjured mice (regardless of sham-status) are grouped together in our analyses (but figures include shaded dots to indicate sham vs. naïve).

### Head-post surgery

To reduce motion and susceptibility artifacts during MRI with low-dose (0.5% isoflurane) anesthesia, all mice undergo a minimally invasive surgery where a custom head-plate is affixed to the skull as we have described previously^41–43^. Mice undergo head-plate implantation following the last day of rmTBI/Stress. Briefly, mice are initially anesthetized with 5% isoflurane (70/30 medical air/O_2_) and secured in a stereotaxic frame (KOPF) using ear bars and an incisor bar (∼ 3 minutes). Isoflurane is then reduced to 2%. Ointment is applied to the eyes to prevent dryness; meloxicam (2mg/kg body weight) is administered subcutaneously, and bupivacaine (0.1%) injected locally at the incision site. After hair removal (Nair, chemical hair remover), the scalp is washed 3 times with betadine followed by ethanol 70%. The skin and soft tissue overlying the skull is surgically removed. The skull is cleaned and dried. Antibiotic powder (Neo-Predef) is applied to the incision site, and isoflurane reduced further to 1.5% (∼ 10 minutes). Superglue (Locite) is applied to the exposed skull, followed by transparent dental cement C&B Metabond (Parkell). The pre-cut head-plate is attached to the dental cement prior to its solidification. The entire procedure is concluded in ∼ 20 minutes, recovery takes < 10 minutes.

### PET data acquisition, processing, and analyses

#### Data acquisition and processing

All mice undergo [^18^F]SynVesT-1 PET scans after completing treatment, or 6-weeks after the last TBI (**Figure 1**), on a Focus 220 scanner (Siemens Medical Solutions). A ^57^Co transmission scan is collected for attenuation correction. During acquisition, mice are maintained on isoflurane anesthesia (1.5% – 2.5%). Three mice are placed in the scanner during each PET acquisition. [^18^F]SynVesT-1 is administered via intramuscular injection (< 0.2 mL). The injected radioactive doses were 25.0 ± 10.1 MBq. After acquisition, anesthesia is discontinued, and recovery monitored closely (mice recover in < 10 minutes). Thus, the entire imaging procedure lasts < 90 minutes.

Images are reconstructed with a 3D ordered subset expectation maximization method (3D-OSEM; 2 iterations, 9 subsets) with maximum *a posteriori* probability algorithm (MAP; 18 iterations) with corrections for decay, random effects, attenuation, and scatter. Images of SUVs (i.e., activity normalized to injected dose and body weight, or voxel value in the reconstructed image (Bq/mL) • body weight (g)/injected dose in Bq) are generated using data from 30-60 minutes post-injection. SUVRs are normalized to striatum. This region was selected based on the effect size for group differences (controls vs. injured) was the lowest in this region. One of the major challenges in rodent PET studies is quantification. In this study, we utilized the averaged SUV from 30-60 minutes normalized by the striatum. The optimal reference region in this study would be an area unaffected by CVS or rmTBI. Although previous literature indicates that TBI might induce presynaptic terminal degeneration in the striatum^44^, our findings show that the striatum SUV (normalized by injected activity and body weight only) exhibited the smallest group differences on average and lowest effect size. Based on those findings, we hypothesize that normalization by striatum values has the smallest impact on the group differences while minimizing the data variability.

#### Analyses

FSL GLM is used to perform a statistical comparison of SUVRs across the entire brain between injured and control mice. We perform nonparametric permutation testing with 5,000 permutations, using family-wise error correction with TFCE. Statistical significance is defined as *p* < 0.05. Based on the significant results obtained from this group comparison, we create a group of ROIs (**Figure 3B**). Specifically, we define regions as the intersect between voxels we determine to show synapse losses (SV2A-PET) and regions within the Allen Brain atlas^45^. The cerebellum, olfactory bulb and medulla are excluded from fMRI analyses based on limited coverage. This approach results in 10 regions, which we use in all other analyses (unless specified).

### MRI data acquisition, processing, and analyses

#### Data acquisition

Data are acquired on an 11.7T system (Bruker, Billerica, MA), using ParaVision 6.0.1 software. Body temperature is continuously monitored (Neoptix fiber) and maintained with a circulating water bath at 36.6- 37°C. BOLD-fMRI data are acquired using a gradient-echo, echo-planar-imaging (EPI) sequence with a repetition time (TR) of 1.8 seconds, and echo time (TE) of 9.1 ms. Data are isotropic 0.275 × 0.275 × 0.275 mm^3^, with 35 slices yielding near whole-brain coverage (no gaps). Each functional imaging run is comprised of 334 repetitions (∼10-minutes). In total, 30-minutes of resting-fMRI data are acquired during each imaging session (i.e., from each mouse).

##### Structural MRI

These data are used for assessment of physical damage (**Figure 2**) and registration to a common space as we have described previously^41, 43^. [**1**] A high in-plane resolution image of the fMRI field-of-view. Using a MSME sequence, in 7min 20s, using a TR/TE of 2500/20ms, we obtain 35 slices (0.275 mm thick) with an in-plane resolution of 0.1 × 0.1 mm^2^. The slice prescription matches those of the fMRI data (i.e., they are of the same anatomy). [**2**] Isotropic 3D anatomy of the whole brain. Using a MSME imaging sequence, in 5min 20s, using a TR/TE of 5500/15ms, we obtained a 0.2 × 0.2 × 0.2 mm^3^ (single average) image of the whole brain.

**Figure 2:**
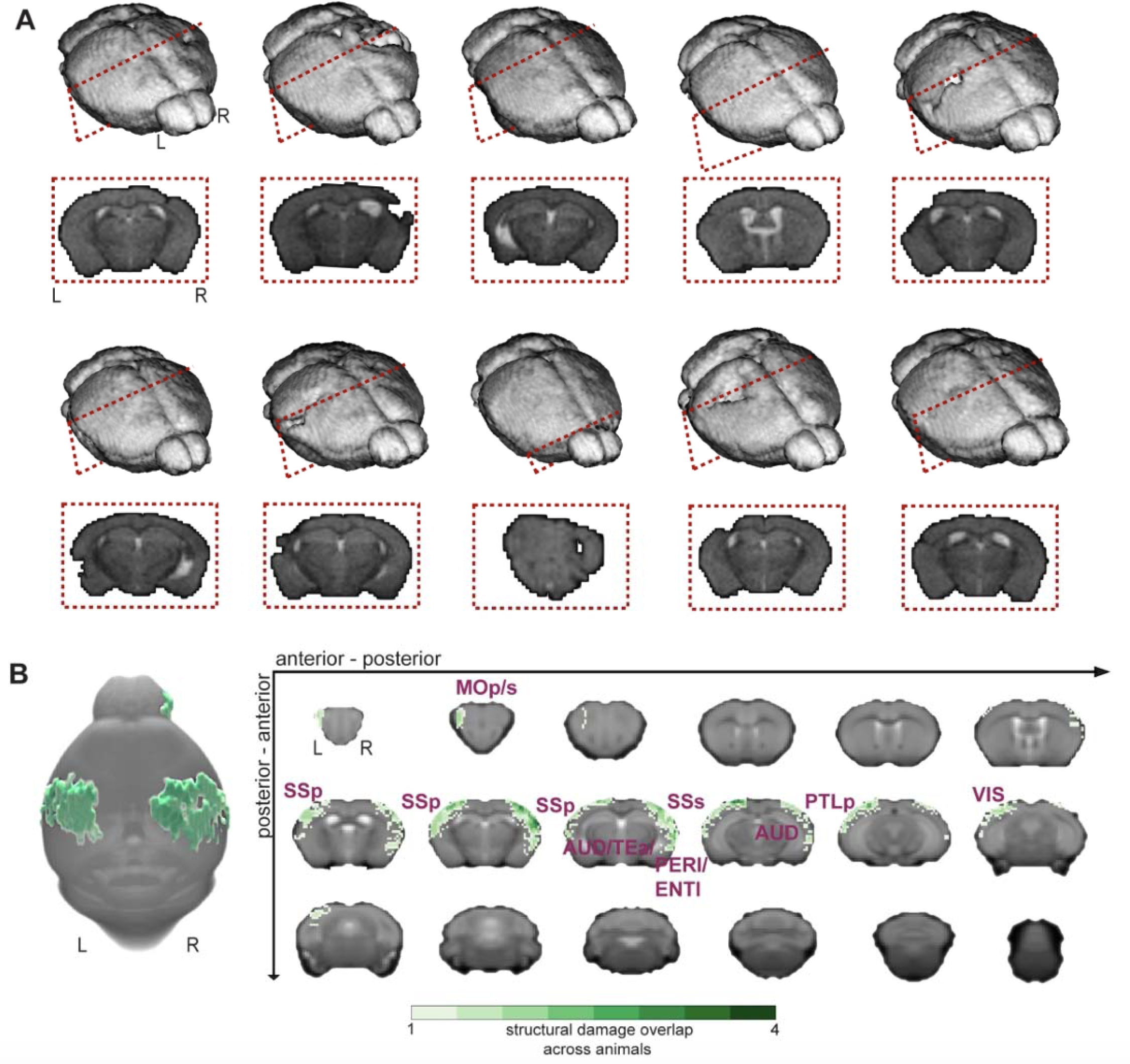
Physical damage (edema) from repetitive mTBI/stress. **A)** 3D reconstructions (BIS, BioImage Suite) of each mouse brain in the injured group (one image, or imaging slice, per mouse). Coronal images displaying the area with the most damage appear below each 3D image with hyper-intense voxels removed or masked-out. Injuries are mild and concentrated in cortical areas at or near the impact sites. **B)** Physical damage across all (N=10) mice. Overall, the group is heterogeneous, with a maximum of 4 animals showing physical damage in the same location, and the physical injuries are mild. The color bar indicated voxels which show damage across all mice in the injured group (white – corresponds to damage appearing in 1/10 mice, green – corresponds to damage appearing in 2-4/10). The affected cortical areas include MOp/s – motor area primary/supplementary, SSp/s – Somatosensory area primary/supplementary, AUD – auditory area, PTLp – posterior parietal association areas, including TEa – temporal association area, VIS – visual areas, PERI/ENTI – perirhinal/entorhinal lateral areas.

After structural and functional MRI data have been acquired, anesthesia is discontinued, and recovery monitored closely (mice recover in < 10 minutes). Thus, the entire imaging procedure lasts < 60 minutes.

#### Data processing

Functional MRI data are processed using the open-source RABIES (Rodent Automated BOLD Improvement of EPI sequences) software package (https://github.com/CoBrALab/RABIES)^46^. A volumetric EPI is derived from trimmed average EPI frames after motion realignment. Using this volumetric EPI as a target, the head motion parameters are estimated by realigning each EPI frame to the target using a rigid registration. To conduct common space alignment, structural images, which are acquired alongside the EPI scans, are initially corrected for inhomogeneities and then registered together to allow the alignment of different MRI acquisitions. This registration is conducted by generating an unbiased data-driven template through the iterative nonlinear registration of each image to the dataset consensus average. The average is updated at each iteration to provide an increasingly representative dataset template^47^. The finalized template, after the last iteration, provides a representative alignment of each MRI session to a template that shares the acquisition properties of the dataset (e.g. brain shape, FOV, and anatomical contrast), which makes it a stable registration target for cross-subject alignment. After aligning the MRI sessions, this newly-generated unbiased template is then itself registered, using a nonlinear registration, to an external reference atlas to provide both an anatomical segmentation and a common space comparable across studies defined from the provided reference atlas. To correct for EPI susceptibility distortions, the volumetric EPI is also subjected to inhomogeneity correction, and then registered using a nonlinear registration to the anatomical scan from the same MRI session, which allows the calculation of the required geometrical transformations for recovering brain anatomy^48^. Finally, after calculating the transformations required to correct head motion and susceptibility distortions, slice timing correction is applied to the timeseries (3dTshift)^49^. Preprocessed timeseries in common space are generated by concatenating the transforms allowing resampling to the reference atlas (the Allen Brain Atlas)^45^, at a voxel resolution of 0.2 x 0.2 x 0.2 mm^3^.

Confound correction of EPI timeseries data is conducted in common space. First, frames with prominent corruption are censored (i.e. scrubbing, see Power et al. 2012)^50^. Framewise displacement^50^ is measured across time and each frame surpassing 0.075mm of motion, together with 1 backward and 2 forward frames, are removed. Next, voxelwise detrending is applied to remove first-order drifts and the average image. Selected nuisance regressors are then used for confound regression. More specifically, using ordinary least square regression, the 6 rigid motion parameters, the mean signal from the cerebral spinal fluid mask and the global signal are modelled at each voxel and regressed from the data. Finally, a spatial Gaussian smoothing filter(nilearn.image.smooth_img)^50, 51^ is applied at 0.3mm full-width at half maximum (FWHM).

#### Analyses

##### Physical damage

Lesion volume is assessed by manual segmentation of MSME (multi-spin, multi-echo) MRI data. Voxels with signal intensities above or below average grey/white matter are delineated in native space using tools developed for this purpose within BioImage Suite (BIS, https://bioimagesuiteweb.github.io/alphaapp/index.html).

##### ReHo

Voxelwise ReHo maps are computed using AFNI^49, 52^. ReHo is calculated for a given voxel and its 27 nearest neighbors and smoothed by a Gaussian kernel with FWHM equal to 10.

##### fALFF

Voxelwise fractional Amplitude of Low Frequency Fluctuations (fALFF) is assessed as a ratio of BOLD-fMRI activity in the low frequency range (0.01-0.075Hz) to the entire frequency range (0.01-0.03Hz)^53, 54^.

##### FC strength

Correlation matrices are obtained using the whole-brain Allen atlas (**Figure S2**) or ROIs generated from SV2A-PET data (**Figure 3B**). Pearson’s correlation is computed between all region pairs.

**Figure 3:**
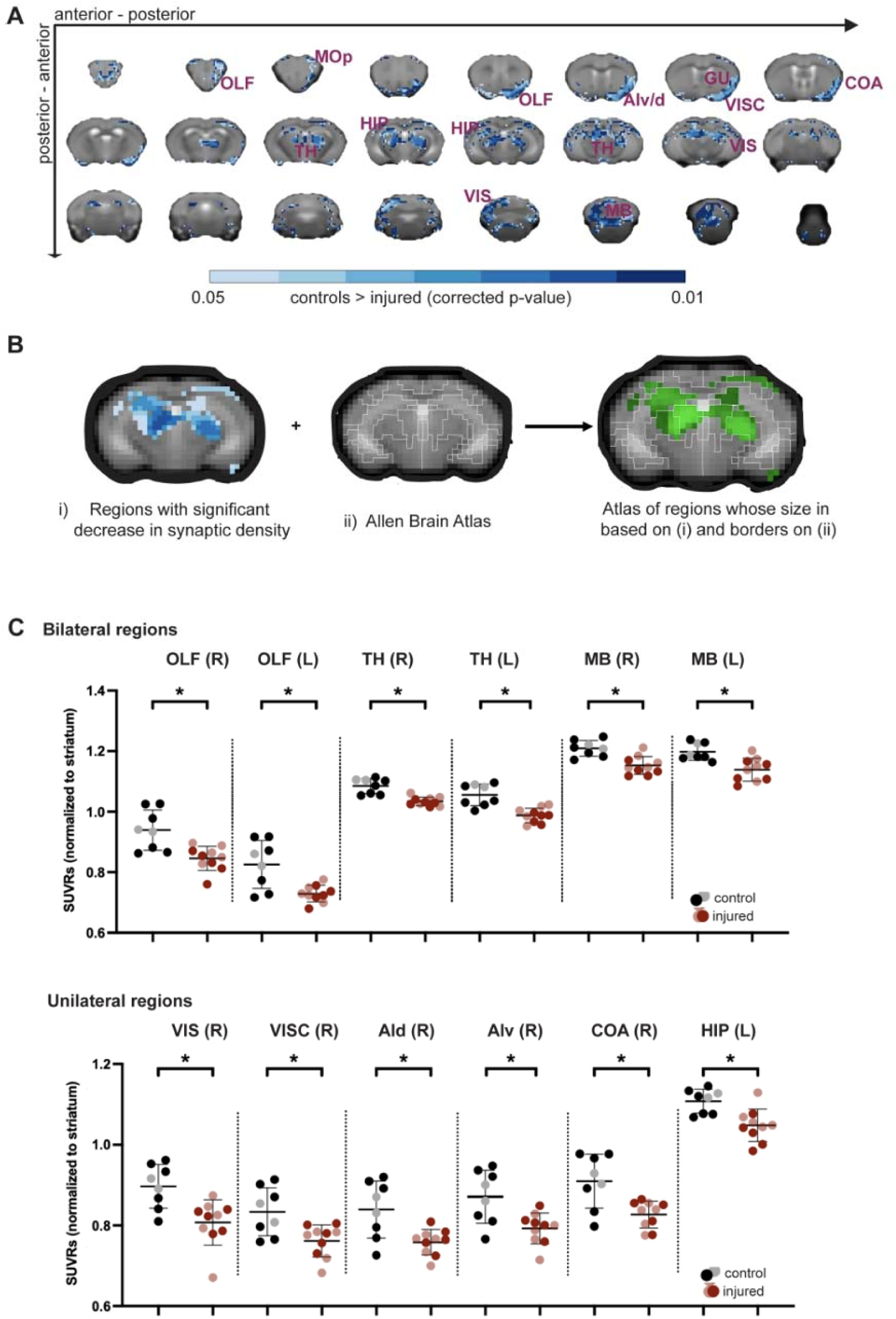
Widespread decreases in synapse density in ‘spared’ brain areas. **A)** In injured mice, regions across the brain show decreases in synapse density compared to controls (threshold-free cluster enhancement – corrected). Affected areas include: OLF – olfactory areas, MOp – primary motor area, VIS - visual area, VISC – visceral area, GU – gustatory area, Ald/v – agranular insular dorsal/ventral areas, COA – cortical amygdalar area, HIP – hippocampus, TH – thalamus, and MB – midbrain. **B)** We take the intersect of voxels showing a decrease in synapse density and regions defined by the Allen Brain Atlas ^41^. Areas showing physical damage (Figure 2) are excluded. **C)** Standardized uptake value ratios from each mouse are plotted. An independent t-test (two-tailed, false discovery rate corrected, p<0.005) indicates significant decreases in synapse density in injured relative to control mice. L – left, R – right. Grey circles (control group) indicate data points from sham-mice whereas black circles (control group) indicate naïve controls (see **Methods**). Similarly, dark-red circles (injured group) indicate data from treatment-naïve mice whereas light-red circles (injured group) indicate data from mice in a pilot-treatment group (see **Methods**).

##### Statistics

FSL GLM (FMRIB Software Library)^55^ is used to perform statistical comparison analyses between injured and uninjured mice. We perform nonparametric permutation testing with 5,000 permutations, using familywise error correction with TFCE. Statistical significance is defined as *p* < 0.05. Additionally, independent t-tests are used to ascertain differences between groups. Specifically, average SUVRs (**Figure 3C**), ReHo (**Figure 4C**) and fALFF (**Figure S5**), which are adjusted using FDR corrections for multiple comparisons^56^. Data and code are available upon request.

**Figure 4:**
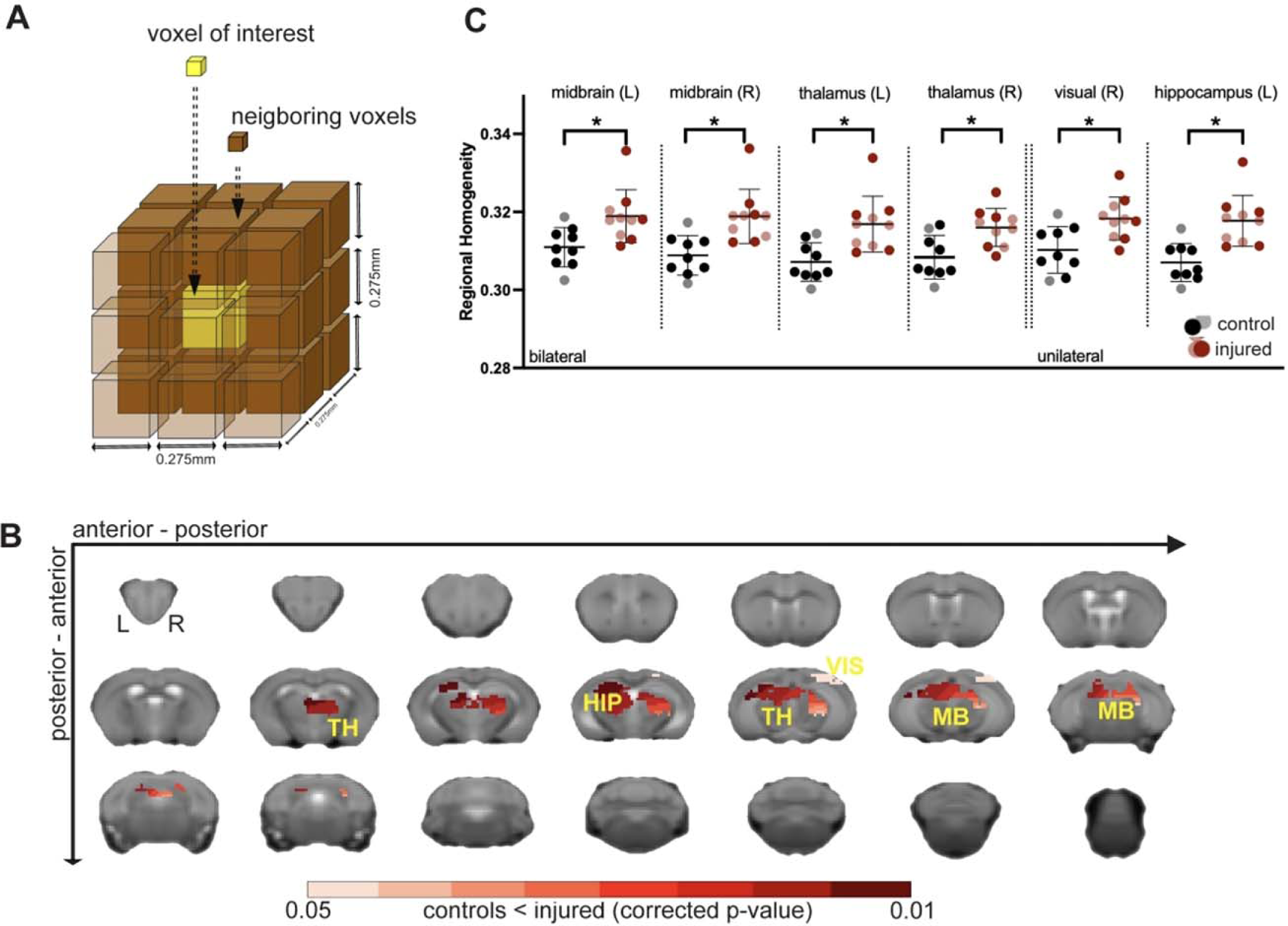
Increased BOLD-fMRI ReHo in regions with depleted synapse densities. **A)** A graphical representation of the BOLD-fMRI ReHo metric ^48^. **B)** Generalized linear model analyses indicate an increase in ReHo in injured mice relative to controls (threshold-free cluster enhancement – corrected) within areas showing a decrease in synapse density (Figures 3) and are removed from the sites of physical damage (Figures 2). **C)** Regional homogeneity (ReHo) from each mouse is plotted. An independent t-test (two-tailed, false discovery rate corrected, p<0.005) indicates a s gnificant increase in ReHo in injured animals compared to controls. L – left, R – right. Color-coding is as in Figure 3.

## Results

We used daily repeated mTBI in combination with stress exposure over 14 days weeks as a mouse model of combined brain injury and chronic stress (**Figure 1**)^16^. Six-weeks after injury, mice underwent two scan sessions, one using the PET tracer, [^18^F]SynVesT-1, a ligand of SV2A as a pan-neural synapse density marker^57–63^, and a second scan session using MRI (**Figure 1**). Our goal was to investigate the emergent functional abnormalities that follow injury and stress using clinically accessible *in vivo* imaging modalities – and the intermodal relationship of their complementary contrasts.

### Mild and heterogeneous tissue damage or edema in cortical areas at or near the impact sites

Our goal was to induce a mild injury with limited destruction of brain tissue. To assess our success, we quantified changes in T_2_-contrast using MSME-MRI (**Methods**). Voxels with high intensities (likely indicative of edema^64^) were segmented manually by a blinded observer (no hypo-intensities were noted). **Figure 2A** displays 3D reconstructions (BioImage Suite, BIS, https://bioimagesuiteweb.github.io/alphaapp/index.html) of each injured brain. An example coronal image, from each mouse highlights the area where the most evidence of damage was observed. Overall, these data indicated minimal destruction of brain tissue, and an appreciable amount of heterogeneity within the group – akin to the human TBI patient population. **Figure 2B** summarizes these data. A maximum of four out of 10 mice had overlapping physical damage (i.e., the same voxels were identified across mice), confirming the heterogeneity within the dataset and the mild nature of the induced injuries (on average). Mostly cortical regions at or proximal to the impact sites were affected (as expected).

### A widespread decrease in synapse density followed injury in ‘spared’ brain areas

Six-weeks after injury (**Figure 1**), we assessed changes in synapse density via SV2A-PET. Specifically, we used a PET tracer, [^18^F]SynVesT-1, which targets SV2A (**Methods**). This marker correlated well with *in vitro* histopathological synapse markers and thus allowed for an *in vivo* evaluation of synapse density^57–63^. PET Standardized uptake value ratios (SUVRs), using striatum as the reference region, from injured and control mice were compared at the voxel-level (FSL General Linear Model, GLM). **Figure 3A** shows widespread decreases in synapse density in regions that showed no physical damage (on MRI) in the injured relative to the control group (p<0.005; threshold free cluster enhancement, TFCE; corrected; uncorrected results are shown in **Figure S3**). Based on the intersection of affected voxels and the Allen Brain Atlas ontology^45^ (**Figure 3B**), we compared average SUVRs from each mouse (**Figure 3C**, independent t-test two-tailed, FDR corrected, p<0.005). Reduced synapse density was observed bilaterally in olfactory areas, thalamus, and midbrain. Unilateral decrements were more prominent on the right side and observed in visual and visceral areas, hippocampus, agranular insular dorsal/ventral areas, and the cortical amygdala area.

### Increased BOLD-fMRI Regional Homogeneity (ReHo) in areas with depleted synapse densities

We assessed the effects of rmTBI/Stress on fMRI measures of ReHo, a metric of local BOLD synchrony (correlation strength among neighboring voxels)^52^. Specifically, within regions with synapse loss (as defined in **Figure 3B**), ReHo was computed for each voxel and its 27 neighbors (**Figure 4A**), and compared between injured and control groups (FSL, GLM). The olfactory bulb was excluded due to a lack of coverage in fMRI data. Significant but uncorrected changes in ReHo are shown in **Figure S4**. Across injured mice, a significant (p<0.05, corrected) increase in ReHo was observed relative to the control group within the thalamus, midbrain, left hippocampus and right visual area (**Figures 4B-C**).

To assess changes in regional excitability, we computed voxelwise the fractional Amplitude of Low Frequency Fluctuations (fALFF)^53, 54^ as the ratio between BOLD activity measured in the low frequency (0.01-0.075Hz) and entire frequency (0.01-0.03Hz) range (**Figure S5**). This metric is thought to capture neural excitability^53, 54, 65, 66^ and was thus one way to assess regional (hyper)excitability. Our results did not show a significant change in fALFF, although a statistically non-significant trend towards an increase in fALFF was observed in the hippocampus and thalamus (p<0.05, uncorrected) of injured mice (**Figure S5**).

### In injured animals, regions with higher synapse densities showed a greater increase in local BOLD-fMRI synchrony

Our multimodal data offered us the unique opportunity to examine inter-contrast relationships. We first assessed how synapse density (SUVR) and regional homogeneity (ReHo) were related in control mice, i.e., without any exposure to rmTBI/Stress. Average SUVR and ReHo were extracted from regions following the Allen Brain Atlas^45^. Results indicated a moderate but significant positive correlation (Pearson’s rho =0.582; p<0.001) between ReHo and SUVR in the healthy condition (**Figure 5A**). This means that the regions with higher standard uptake values (SUVR) showed greater regional homogeneity (ReHo).

**Figure 5:**
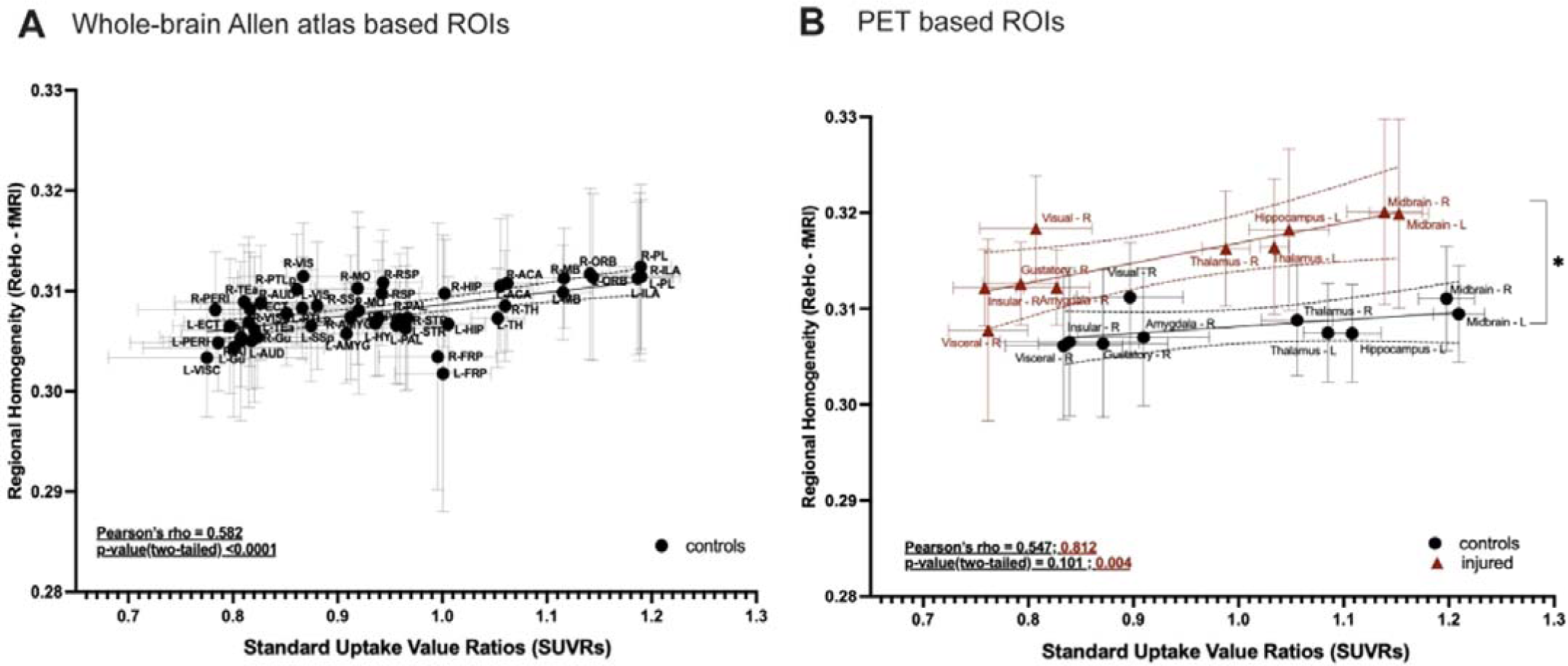
Relationship between synapse density (SV2A-PET SUVRs) and ReHo (BOLD-fMRI). **A)** Regions from the Allen Brain Atlas ^41^ are used to calculate regional homogeneity (ReHo) and synapse density in the control group (N=10). Measures are averaged for each region and Pearson’s correlation is applied. A high correlation is uncovered between these PET and fMRI measures. **B)** Similar to A), ReHo and synapse density are computed within regions ‘spared’ from physical damage (Figure 2) that show injury-induced decreases in synapse density (Figure 3B). Measures are averaged for each region for each group and Pearson’s correlation is applied. Data from injured mice appear in red. Data from control mice appear in black. A difference between slopes is found (t-value = 2.15; Dfd=16; p-value, two-tailed=0.047). SUVRs (standardized uptake value ratios).

Next, we assessed the relationship in injured mice (N=10). Average SV2A-PET SUVRs and BOLD-fMRI ReHo values were extracted from N=10 regions (**Methods**) as defined in **Figure 3B**. Between measures, a positive correlation (Pearson’s rho =0.582; p<0.001) was uncovered (**Figure 5A**). This was compared, using the same regions, to the control group. A strong and statistically significant correlation (Pearson’s rho =0.812; p=0.004) was found in the injured group (**Figure 5B**). A statistically insignificant trend towards a moderate positive correlation (Pearson’s rho =0.547; p=0.101) was uncovered in controls (likely due to our small ‘N’ and effect size). Further, a difference between slopes across groups was observed (t-value = 2.15; Dfd=16; p-value, two-tailed=0.047). In summary, SV2A-PET SUVRs, and BOLD-fMRI ReHo showed a positive correlation in healthy mice (across the whole brain). With injury, this relationship (between SV2A-PET SUVRs and BOLD-fMRI ReHo) strengthened in ‘spared’ brain regions. Moreover, separate from the impact foci and in areas without any physical damage on structural MRI, variation in synapse density was highly correlated with an increase in ReHo, such that regions with *higher synapse* densities showed a *greater increase* in ReHo after injury.

### Sparse indications of increases in BOLD-fMRI functional connectivity were uncovered following injury

In addition to assessing local BOLD-fMRI synchrony within regions, we assessed injury-induced changes in inter-regional FC strength^67^. We assessed brain-wide functional connectivity calculated based on the Allen Brain atlas^45^ (**Figure S6A**). Permutation testing (5,000 permutations) indicated no significant (uncorrected) differences between groups. However, among ‘spared’ regions showing decreases in synapse densities (**Figure 3**), FC strength (z-scored) in injured relative to control mice (**Figure 6A**) showed a decrease between the right agranular and visual areas, as well as more widespread increases including between gustatory and visual areas, thalamus and agranular areas and left and right thalamus (FSL, GLM) (**Figure 6B**). These analyses were exploratory (p<0.005, uncorrected), as our observations did not survive FDR correction for multiple comparisons.

**Figure 6:**
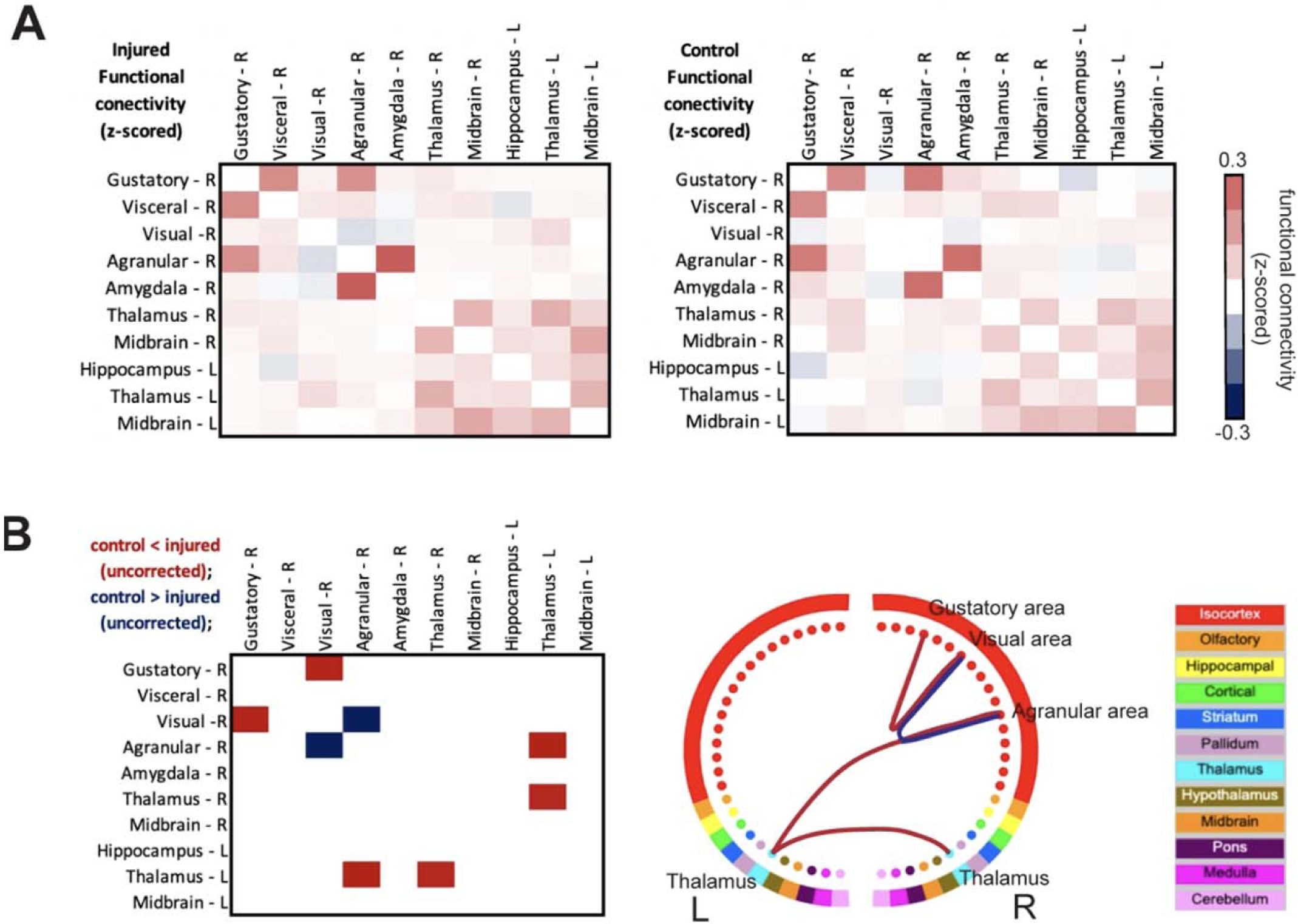
Patterns of injury-induced changes in BOLD-fMRI FC strength. **A)** BOLD-fMRI FC strength (z-scored) between all ‘spared’ region pairs showing decreases in synapse density (Figure 3B) for mice in injured (left) and control (right) groups. **B)** Generalized linear model with permutation testing (5,000 permutations) is used to compare FC strength between groups. Red indicates an increase in FC strength (p<0.005, uncorrected) in injured relative to control mice. Blue indicates a decrease in FC strength (p<0.005, uncorrected) in injured relative to control mice. L – left, R – right. Circle plot generated using BIS.

## Discussion

Repetitive-mild traumatic brain injury (rmTBI) can lead to debilitating protracted neurological and behavioral symptoms that can have a delayed onset. Clinical standardized treatment methods to properly manage patients, and effectively monitor and treat rmTBI symptoms and potential progression are lacking. To address these gaps, preclinical studies can play a crucial translational role. Here, we use a mouse model which combines rmTBI and CVS to recapitulate key phenotypes of rmTBI/Stress. To this end, we and others have shown that stress exacerbates mild mechanical trauma^15, 16^ and subsequent tauopathy^12^.

Six-weeks post-injury, mice undergo PET with an SV2A ligand and MRI. With a focus on translational relevance, we use clinically assessable imaging modalities to characterize injury-induced brain abnormalities. Injured mice show limited and heterogeneous physical damage (MRI), but widespread decreases in synapse density – including in regions ‘spared’ from physical damage (on structural MRI) – coupled with an increase in BOLD-fMRI ReHo. In uninjured mice, a positive correlation is found between SV2A-PET SUVRs and BOLD-fMRI ReHo. This relationship is enhanced following injury, suggesting that brain areas with *higher* synapse densities show a *greater increase* in ReHo post-injury. Additionally, sparse increases in FC strength are observed. Taken together, these results may reflect endogenous functional compensation for injury (more below).

### A widespread decrease in synapse density accompanies an increase in BOLD-fMRI ReHo

SV2A is ubiquitously expressed in presynaptic vesicles, making it a suitable *in vivo* biomarker for synapse density measurements^68, 69^. Recent validation studies in humans^61, 70–73^ and non-human primates^74, 75^ include pioneering work measuring synapse losses in patients with temporal lobe epilepsy^71^, and severe depression^73^. Studies in rodents, validated with *in vitro* immunoreactivity^40, 69^ show synapse losses in a mouse model of AD^57, 58^, and in rat cortical and hippocampal regions following a single head injury^76^. Our findings align well with previous SV2A-PET studies. After rmTBI/Stress, mice show a decrease in synapse density, not only in cortical and hippocampal areas, but also in the cerebellum, midbrain, olfactory areas, and thalamus.

To assess how decreases in synapse density affect local connectivity we measure BOLD-fMRI synchrony using ReHo^52^. In patients with TBI, ReHo, at various times after injury, shows markedly heterogeneous results in terms of the brain regions and circuits affected, as well as whether ReHo goes up or down (**Table 1**)^25–30, 77, 78^. Given the heterogeneity in time after injury, mechanism of injury (e.g., military service or sports-related), and injury location, there is a clear need for further research.

**Table 1.** Summary of BOLD-fMRI ReHo measurements in TBI patients.

| Study | TBI/healthy | Time post-injury | Injury criteria | ReHo | Finding |
| --- | --- | --- | --- | --- | --- |
| Zhan et al., 2015 | 15/15 | < 2 weeks | Single concussion no damage on structural MRI | Decrease | Insula L, precentral/postcentral gyrus L, & supramarginal gyrus L |
| Wu et al., 2017 | 18/18 | Within 6 months | Acceleration-deceleration or high velocity rotation | Decrease | Bilateral thalami (ventral anterior & lateral nuclei) |
| Liu et al., 2020 | 32/37 | Within 7 days | Traffic accident (9), falls (2), sports-related (2), object strike (19) | Increased | Rolandic operculum L, heschl gyrus L, & superior temporal gyrus L |
|  |  |  |  | Decreased | Bilateral calcarine fissure & surrounding cortex, cuneus L, lingual gyrus L, & bilateral thalamus |
| Meier et al., 2017; 2020 | 30 (one day), 34 (one week), 30 (one month), 51 controls | 1 day - 1 month | Sports-related concussion | Increasing over time | Sup motor area, superior temporal gyrus R, bilateral postcentral gyrus, paracentral gyrus L, middle temporal gyrus R, lingual gyrus R, fusiform gyrus R |
|  |  |  |  | Decreasing over time | Superior medial frontal gyrus, mid frontal gyrus R, & superior frontal gyrus R |
|  | 92/82 | 24-48 hours, at 7 days, and 6 months | Sports-related concussion, no damage on structural MRI | Increased | At 24 hours: superior & mid frontal gyri R |
| Shi et al., 2021 | 50/50 | < 3 days | Traffic accident (28), daily living (9), sport (13) | Decreased | Inferior temporal gyrus, cerebellum |
| Vedaei et al., 2021; 2023 | 25/40 | > 6 months since last injury | One or more injuries | Increased | Occipital mid L, & cingulate post L |
|  | 60/40 | 24-37 months | One or more injuries, single: multiple (17:43) | Increased | Occipital mid L, fusiform R, lingual L, & putamen L |
|  |  |  |  | Decreased | Frontal sup L, temporal sup R, frontal sup medial L, & OFC posterior L |

L – left, R – right, mid - middle, Sup – supplementary, OFC – orbital frontal cortex.

Experimental animal models of TBI are likely to improve our understanding of brain injury progression, and its imaging correlates, given our ability to control injury mechanism, location, frequency, and severity. Studies on rodent TBI models measuring BOLD-fMRI ReHo are just beginning to emerge. We identified two in the literature. The first is a longitudinal study of mice following a single CHI^79^. Despite the difference in model (rmTBI/Stress vs. a single CHI), and timepoint (6 weeks vs. 2 days and 2 weeks) our findings align. To et al. observe an increase in ReHo in similar brain regions including the hippocampus, thalamus, and midbrain^79^. On the other hand, a more recent study on drug seeking mice at a later timepoint (7 weeks post-injury), using a different injury mechanism (3 repeated blast injuries), uncovers very sparse (or negligible) increases in ReHo, but more widespread increases when the interaction between injury and drug seeking behavior is considered^80^. Given the potential for translation to the clinic, future work is needed to determine the effects of injury mechanism, and time, on BOLD-fMRI measures of ReHo in animal models of (rm)TBI.

Increases in ReHo could be related to p-tau, neuroinflammation, and/or neural hyperexcitability. Our previous work in a CHI mouse model^16, 18, 24^ uncovered p-tau and neuroinflammation similar to observations made in animal models of AD^12, 58, 81^. In AD, we and others have established a link between p-tau, neuroinflammation, and synapse losses measured via SV2A-PET^58, 72, 82^. P-tau and neural hyperexcitability have also been reported in mouse models of AD^83, 84^ and brain injury^37^, alongside increases in astrocytic activity^85^. In AD, an increase in hippocampal ReHo, a positive correlation between ReHo and tau-immunoreactivity, and evidence of neural hyperexcitability, have been reported^86^. Taken together, accumulating evidence seems to suggest that brain injury or AD can elicit p-tau coupled with an immune response and neural hyperexcitability which in turn manifests as an increase in BOLD-fMRI ReHo. To & Nasrallah proposed that hyperexcitability is responsible for their observations of increased ReHo and BOLD-fMRI FC, in a single concussion TBI model^38^. One interpretation of increased ReHo and neural hyperexcitability is that it reflects evidence of endogenous compensation for injury (or disease) – neurons attempting compensate which could result in greater local synchrony. Future studies are needed to dig deeper into these speculations.

To provide preliminary exploration of the hypothesis that increases in ReHo reflect neural hyperexcitability, we compute voxel wise fALFF^53, 54^ (**Figure S5**). This metric is thought to capture neural excitability^53, 54, 65, 66^ and is thus one way to probe our hypothesis. Encouragingly, TBI patients at three months post-injury, show increases in fALFF within regions that that show increases in ReHo, and there is a significant correlation between the two metrics^30^. However, to the best of our knowledge, fALFF has not yet been reported in an animal study of brain injury. While our results do not show a significant change in fALFF, a statistically non-significant trend towards an increase in fALFF is observed in the hippocampus and thalamus (p<0.05, uncorrected) of injured mice (**Figure S5**). Further studies are warranted to reliably assess changes in fALFF caused by brain injury and to explore relationships between BOLD-fMRI measures (ReHo, FC, and fALFF).

To the best of our knowledge, this is the first study to compare SV2A-PET and BOLD-fMRI measurements in the same brain. Thus, we begin by quantifying the relationship between these metrics in healthy (uninjured) mice. We find a robust positive correlation between SV2A-PET SUVs and BOLD-fMRI ReHo across the whole-brain. This suggests that BOLD-fMRI ReHo is (to some extent) reflective of underlying synapse densities. In brain regions susceptible to injury-induced synapse loss, this correlation strengthens with injury and ReHo changes are greatest in regions with greater synapse density. The brain regions with high excitatory synapse densities include hippocampus, midbrain, cerebellum, and thalamus^87^. The dependence of injury-induced ReHo on baseline synapse density is consistent with the hypothesis of increased post-injury excitability. This hypothesis requires further exploration which takes into account both pre- and post-injury measures.

Although we observe sparse changes in BOLD-fMRI FC strength that do not survive corrections for multiple comparisons, the circuit showing the greatest numerical change aligns well with underlying anatomical connectivity. The gustatory cortex projects to the hippocampus, amygdala, visceral and agranular (insular) cortex, thalamus, and visual cortex. Insular areas project to the amygdala, while agranular areas project to parts of the thalamus and midbrain (**Figure S7**). Collectively this circuit is responsible for touch, taste, and olfaction^88–90^. Encouragingly, a similar pattern of increased FC strength at eight weeks after a single controlled cortical impact has recently been reported^91^. Given the collective role of the injury-affected circuit in sensory processes, it would be intriguing to study further via coupled behavioral and multimodal imaging experiments. For human TBI, BOLD-fMRI measurements of FC strength show pronounced heterogeneity, matching the variability of ReHo (**Table 1**)^25–30, 77, 78^. This likely reflects the diverse nature of clinical (rm)TBI studies. Yet, relative to BOLD-fMRI ReHo, the FC literature is considerably more developed (see reviews, Lunkova et al. 2021; Mayer et al. 2015; Jantzen et al. 2010; Eierud et al. 2014)^31–34^. Of particular relevance to the present study, brain-wide increases in FC strength, alongside increases in ReHo and fALFF have been recently reported^30^.

### Study limitations

Despite employing rigorous statistical approaches to mitigate the impact of a small sample size, our ability to draw definitive conclusions, especially in fALFF and FC analyses, is limited. Our control group is comprised of both sham and naïve mice (**Methods**). Although we took careful steps to ensure that sham and naïve mice show no differences across measures (see **Methods** and **Figure S1**), a group of sham-only mice would have been preferable. A subset of our injured mice received AZD0530 treatment (saracatinib) or a vehicle as part of a pilot study. Despite the potential neuroprotective effects of AZD0530 in a mouse model of AD^40^, we do not observe any differences between treated mice and those given a vehicle (**Figure S2**). Consequently, we consolidate all injured mice into one group. This study assesses mice with clear yet mild physical brain damage (on MRI), while some studies only include animals with no evidence of structural alterations^35, 68^. Our view is that the heterogeneity and presence of structural damage here provide a closer approximation to the human condition^92^. Further, to address potential irregularities, we exclude all areas with evidence of physical damage from our analyses. Finally, this study is conducted on male mice only. Given the known differences between males and females in TBI^39^, future work will include both male and female animals.

### Conclusions

This study utilizes a mouse model of rmTBI/Stress^93^ over a course of two-weeks. Six-weeks post-injury, mice undergo SV2A-PET and MRI imaging sessions. Our focus is on clinically accessible modalities (PET and fMRI) for assessing emergent abnormalities indicative of risk of neurodegeneration. Our findings reveal widespread decreases in synapse density and increases in BOLD-fMRI ReHo in areas without physical damage. A moderate positive correlation is found between SV2A-PET SUVRs and BOLD-fMRI ReHo measures in uninjured mice. In mice with brain injuries, greater increases in BOLD-fMRI ReHo are observed in regions with higher synapse density. Taken together, these observations may reflect compensatory mechanisms following injury. Further multimodal studies are needed to provide deeper insights into these observations.

### Transparency, Rigor and Reproducibility Statement

All experiments and procedures adhere to the protocols approved by the Institutional Animal Care and Use Committee (IACUC - 2023-20284) at Yale University. Twenty-one wildtype male mice are randomly allocated into control and injured groups, guided by prior research from the laboratory, < 10 animals per group is sufficient to detect behavioral and brain changes. Repetitive mild traumatic brain injury (rmTBI) with stress is administered daily in a randomized manner. Specifically, rmTBI comprises a two-week regimen involving daily exposure to three out of five aversive stimuli: (1) a 3-minute cold water swim (16-18°C), (2) overnight food deprivation with access to water ad libitum, (3) 3 hours in a cage with 300ml of water added, (4) 3 hours exposed to a cage tilted at a 45° angle, and (5) 15 minutes of immobilization in a flat-bottomed restraint chamber (Braintree Scientific Instruments). The aversive stimuli are randomized to mimic the unpredictable nature of psychological trauma and to limit habituation. Sham-stressed mice are transferred to a single-occupancy cage during the stressor. Daily rmTBI occurs within 1-3 hours of stress exposure (further details in the Methods under “Repetitive Mild Closed-Head Injury with Chronic Variable Stress”). A subset of five randomly selected mice receive a sub-therapeutic dose of saracatinib treatment daily for six weeks at doses of 5mg/kg/day, administered in two equal aliquots via oral gavage until the study endpoint. Independent t-tests, followed by false discovery rate (FDR) correction, show no significant effect of treatment (**Fig S2**) when comparing treated and untreated injured mice. Consequently, treated and untreated mice are combined and treated as the injured group. Structural damage assessments using MRI scans are conducted by a blinded experimenter. Cortical regions showing structural damage are excluded from further analysis, and PET and fMRI data are preprocessed according to standardized protocols and analyzed as outlined in the Methods sections (“PET/MRI Data Acquisition, Processing, and Analyses”). Statistical comparisons between control and injured groups are conducted using nonparametric testing with 5000 permutations, corrected using family-wise error correction with threshold-free cluster enhancement (TFCE), setting statistical significance at p<0.05, unless otherwise specified. Replication studies involving both sexes are currently being planned, and efforts to secure funding are ongoing.

## Acknowledgements

M.M. is funded by SNSF Postdoc.Mobility grant (214392). The authors thank the Yale PET Center staff for their professional technical assistance on the PET tracer production and animal PET scans. Z.C. is funded by NIH grants R01NS123183, R01AG058773, and RF1NS130069.

## Author contributions

M.M – Conceptualization, Data curation, Formal analysis, Visualization, Writing – original draft, review and editing. F.M. Conceptualization, Methodology, Data curation, Data acquisition, Writing – review and editing. T.T. – Conceptualization, Methodology, Data acquisition, Formal analysis, Writing – review and editing. J.C. – Supervision, Funding acquisition, Writing – review and editing. A. F-Z. – Conceptualization, Methodology, Data acquisition, Writing – review and editing, X.S. – Formal analysis, Writing – review and editing. S.M.S. – Conceptualization, Methodology, Formal analysis, Supervision, Funding acquisition, Writing – review and editing. E.L. – Conceptualization, Methodology, Data curation, Formal analysis, Supervision, Funding acquisition, Writing – review and editing.

